# Spring frost controls spring tree phenology along elevational gradients on the southeastern Tibetan Plateau

**DOI:** 10.1101/158733

**Authors:** Yafeng Wang, Bradley Case, Sergio Rossi, Liping Zhu, Eryuan Liang, Aaron M. Ellison

## Abstract

Temperature is considered to be a main driver of spring phenology, whereas the role of climate extremes (such as spring frosts) has long been neglected. A large elevational gradient of mature forests on the Tibetan Plateau provides a powerful space-for-time ‘natural experiment’ to explore driving forces of spring phenology. Combining 5-yr of in situ phenological observations of Smith fir (*Abies georgei* var. *smithii*) with concurrent air temperature data along two altitudinal gradients on the southeastern Tibetan Plateau, we tested the hypothesis that spring frost was a major factor regulating the timing of spring phenology. Onset of bud swelling and leaf unfolding in the study years occurred ≈ 18 or 17 days earlier, respectively, at the lowest (3800 m a.s.l.) elevation relative to upper treelines (4360 or 4380 m a.s.l.). The frequency of freezing events and last freezing date were critical factors in determining the timing of bud swelling along two altitudinal gradients, whereas onset of leaf unfolding was primarily controlled by the onset of bud swelling. This finding provides evidence for detrimental impacts of spring frost on spring phenology, which have been underappreciated in research on phenological sensitivity to climate but should be included in phenology models. It contributes to explain the declining global warming effects on spring phenophases, because climatic extreme events (e.g. spring frosts) tend to increase with warming.

## Introduction

Phenology determines plant survival, growth, and distribution and plays an important role in ecosystem functioning and in the provision of ecosystem services (Chuine & Beaubien, 2001; Inouye, 2008; Forrest & Miller-Rushing, 2010). However, there are uncertainties about the drivers of phenology due to a paucity of a range of intensively-monitored sites (Richardson *et al*., 2013; Gallinat *et al*., 2015; Piao *et al*., 2015). Researchers attempting to generate time series long enough to make inferences about climate-driven changes in phenology often accumulate just one data point per year (Miller-Rushing *et al*., 2010). Furthermore, most tree phenology data are limited to first flowering and leaf unfolding and rarely consider variation in other aspects of tree phenology (e.g., bud swelling) that may respond to climate differently from flowering or leaf unfolding ( Miller-Rushing & Primack, 2008).

For many species in temperate and cold ecosystems, temperature is the key factor controlling onset of spring phenology (Menzel, 2003; Piao *et al*., 2011, 2015; Richardson *et al*., 2013; Huang *et al*., 2014; Laube *et al*., 2014; Chen *et al*., 2015; Davis *et al*., 2015; Fu *et al*., 2015; Ge *et al*., 2015), although other climatic factors (e.g., precipitation, radiation, and photoperiod) can also play a role (Fu *et al*., 2014a, b; Davis *et al*., 2015; Ren *et al*., 2015; Shen *et al*., 2015a). Climate extremes, in particular, are important for understanding the climatic limits of tree species (Inouye, 2008; Zimmermann *et al*., 2009), but few studies have explicitly linked spring phenophases to climatic extremes, such as late-season freezing temperatures (Inouye *et al*., 2008; Ernakovich *et al*., 2014). In spring, plants experience a de-hardening period, during which a certain amount of heat is required to initialize leaf unfolding (Richardson *et al*., 2013; Fu *et al*., 2014b); during this period, plants are particularly vulnerable to freezing events (Lenz *et al*., 2013). Spring freezing events tend to increase in response to climate warming (Inouye, 2000; Augspurger, 2013; IPCC, 2013), and thus late-spring frosts may play an increasingly critical role in controlling spring phenology in temperate and cold regions (Inouye, 2008; Gu *et al*., 2008; Wang *et al*., 2014). Indeed, one recent study showed that spring frost affects the timing of bud-break which, in turn, determines the elevational and latitudinal limits of deciduous broad-leaf tree species in the Alps (Kollas *et al*., 2014). However, less is known about the impacts of spring frost on spring phenology for conifers, which are the dominant species of many subalpine communities.

The Tibetan Plateau hosts mature, natural forests across a broad elevational gradient, which can be used as a space-for-time substitution for longer-term time series data in exploring key drivers of tree phenology. Satellite-based observations and process-based tree-ring growth model have confirmed that advancement of deciduous or semi-deciduous vegetation green-up-date on the Tibetan Plateau over the past three decades is tied closely to spring warming (Piao *et al*., 2011; Zhang *et al*., 2013; Shen et al., 2015b; Yang et al. 2017). However, the impact of spring freezing events on tree phenology is unknown, although spring frosts and freezes occur frequently on the Tibetan Plateau (Shen *et al*., 2014).

We used a five-year dataset of precise, bud-scale measurements of spring phenophase timings to explore climatic drivers of spring phenology for Smith fir (*Abies georgei* var. *smithii*) along two altitudinal gradients. Specifically, we (1) revealed temporal patterns of spring phenophases along two altitudinal gradients; and (2) assessed effects of freezing events and growing degree days on spring phenophases. Given the frequent occurrence of spring freezing events on the southeastern Tibetan Plateau during the period when bud swelling is occurring (Shen *et al*., 2014), we hypothesized that spring frost would be closely associated with the timing of bud swelling.

## Materials and methods

### Study region and climate

The study region is situated in the Sygera Mountains (29° 10’ – 30° 15’ N, 93° 12’ – 95° 35’E) on the southeastern Tibetan Plateau. The south Asian monsoon approaches the Sygera Mountains through the valley of Yarlung Zangbo River, resulting in plentiful summer rainfall (Liang *et al*., 2010). Records from the Nyingchi weather station (29° 34’ N, 94° 28’ E, 3000 m a.s.l.) showed that the mean annual precipitation from 1960 to 2013 was 672 mm, 72% of which occurred from June to September (Liang *et al*., 2010). July (mean temperature of 15.9 °C) and January (0.6 °C) were the warmest and coldest months, respectively.

Based on an automatic weather station (4390 m a.s.l.) near the treeline on the eastern-facing slopes installed in November 2006, the annual average precipitation at our study sites from 2007 to 2013 was 957 mm, 62 % of which fell during the monsoon season (June to September). The warmest and coldest months were July (7.9 ± 0.5 °C) and February (“8.0 ± 1.7 °C), respectively. Snowfall of 50-100 cm usually occurs from November to mid-May.

### Study species and study sites

Smith fir (*Abies georgei* var. *smithii*) is an evergreen coniferous tree species distributed on the north- or southeast-facing slopes on the southeastern Tibetan Plateau (Liang *et al*., 2011). It grows along the altitudinal gradient ranging from 3550 to 4400 m a.s.l., with growth primarily constrained by low temperatures (Liang *et al*., 2010; Li *et al*., 2013, 2017). We studied Smith fir at eight sites along two altitudinal transects: four sites on a southeast-facing slope (labelled as SE3800, SE4000, SE4200 and SE4360, with the number indicating m a.s.l) and four sites on a north-facing slope (N3800, N4000, N4200, N4380) (Fig. S1). At each site, 10 trees were selected for measurement, except for at site SE4200 where only 6 trees were measured. We used a measuring tape to determine the height of each sampled tree in April 2012, when buds were dormant. We estimated tree age by counting the internodes along the main stem (see also the methods in Liang *et al*., 2011).

### Phenology measurements

We made phenological observations on trees weekly between May and September during five consecutive years (2012–2016). From observations of terminal buds we recorded the dates of bud swelling, and leaf unfolding. The onset of bud swelling and leaf unfolding was determined as the dates when trees showed swollen buds or unfolded needles in shoot apices. For all monitored trees, high resolution photographs were taken at each phenological measurement visit with a steel ruler (accuracy of 1 mm) placed behind the shoot apex. In the results, we report “dates” as days elapsed since 1 May each year

### Temperature data

Air temperature (± 0.2 °C) in each stand was measured hourly with a temperature logger (TidbiT v2 Temp UTBI-001, Onset Computer Corporation, Bourne, MA, USA) that was placed 2 m above the ground under the canopy of a tall mature tree. Loggers were not placed under studied trees, which had small main stems. An epoxy radiation shield designed by the logger manufacturer was covered each sensor to minimize effects of direct sunlight on the measurements.

The frequency of spring freezing events was calculated as the number of days with daily minimum temperature < 0 °C in spring (Shen *et al*., 2014). The last freezing date was defined as the last spring day when daily minimum temperature < 0 °C (Schwartz *et al*., 2006). Safety margins were defined as the number of days between the last freezing date and the onset of bud swelling (Dantec *et al*., 2015).

A winter chilling requirement is considered to be an important factor that determines the onset of spring phenophases (Fu *et al*., 2015). We first estimated the number of chilling days as the sum of days in the winter when daily temperature < 0 °C (Yu *et al*., 2010; Fu *et al*., 2015). Accumulated daily mean temperature above a certain threshold, *i.e*., growing degree-days, also has been considered to be an important factor driving the onset of leaf phenology (Fu *et al*., 2014b). As compared to warmer areas, vegetation in colder environment such as Tibetan Plateau requires lower threshold temperature to green up (Piao *et al*., 2011). Thus a minimum temperature of 0 °C was used as a basis to accumulate degree-days, starting with the date when the mean daily air temperature was > 0 °C for at least 5 consecutive days from March and continuing until the onset of bud swelling and leaf unfolding.

### Data analysis

We used regression tree modelling to investigate the influence of the many explanatory factors hypothesized to be important in controlling timing of bud swelling and leaf unfolding (Table 1). For the onset of bud swelling, we investigated the effects of both warmth-related variables (accumulated growing degree-days, minimum, and maximum temperatures) and frost-related variables (frequency of spring freezing events and last spring freezing date). We evaluated the impacts of three variables (elevation, aspect, and year) on the safety margins. For leaf unfolding, we sought to understand how different measures of spring warmth controlled the onset of this phenophase; as leaf unfolding occurred in June, we did not include spring frost variables in these models because frost was not present at this time of the year. In all models, both aspect and year of measurement also were included to control for the effects of these factors related to sampling design (Table 1). As elevation is a proxy for, and was significantly correlated with, several temperature variables in this study, it was not included in the modelling.

**Table 1.** Description ofpredictor variables used in the regression tree modelling of spring phenology (date of onset of bud swelling and leaf unfolding). All climate-related variables were computedfor each site and measurement year based on temperature logger data. BS and LF represented bud swelling and leaf unfolding, respectively.

| Variable name | Description | In Modeling |  |
| --- | --- | --- | --- |
|  |  | BS | LF |
| Degree_days | Cumulative temperatures (above 0°C) from March up to phenophase onset | Yes | Yes |
| Min_tem | Minimum daily temperature recorded at time of phenophase onset | Yes | Yes |
| Max_tem | Maximum daily temperature recorded at time of phenophase onset in each year | Yes | Yes |
| FFE | Number of days/year with daily minimum temperature < 0°C from March to May | Yes | No |
| LFD | Last freezing date in spring (March to May) | Yes | No |
| Aspect | The aspect of a given sample plot | Yes | Yes |
| Year | Year of measurement (2012-2016) | Yes | Yes |
| Chilling days | Number of days with daily temperature < 0°C in winter | Yes | No |

We used a two-stage modelling approach. First we used random forest analysis (Breiman, 2001) to estimate and rank the importance of each explanatory factor in describing variability in the response variables. Second, we used conditional inference trees to gain further insights into the nature of relationships between each response variable and the most important explanatory factors. These two non-parametric, machine-learning methods of analysis allow for the construction of complex, non-linear models with inter-correlated predictor variables (De’ath & Fabricius, 2000; Cutler *et al*., 2007). The random forest approach averages the outcomes of thousands of boostrapped regression trees (‘forests’) to identify those measured explanatory variables that are the best predictor variables. We used the random forest ‘variable importance’ measure to identify the most influential factors in explaining variation in the response variable and then used partial dependence plots to show the marginal effect of each of these factors (*i.e*., while holding all of the other explanatory factors at their average values) on the response variable (Cutler *et al*., 2007). The relative importance of the top-ranked predictor variables was investigated further using conditional inference trees (Hothorn *et al*., 2006) derived from a recursive partitioning method that generates a set of decision rules describing how variation in the response data is best attributed to each predictos. The conditional inference tree method requires a statistically significant difference (*P* < 0.05), as determined by Monte Carlo simulation, to create a partition in the data; this algorithm minimizes bias and prevents over-fitting and the need for tree pruning (Hothorn *et al*., 2006). Random forest and conditional inference tree analyses were implemented in R version 3.1.0 using the ‘randomForest’ (Liaw & Wiener, 2002) and ‘party’ (Hothorn *et al*., 2006) packages, respectively.

## Results

### Tree size

On the southeast-facing slope, trees ranged from 0.47 – 1.57 m in height and were estimated to be 13 – 40 years old (Table S1). The trees on the north-facing slope were 0.39 – 1.83 m in height and 10 – 44 years old (Table S1).

### Among-site variation in phenology

Bud swelling and leaf unfolding began in early May and ended in late May, and occurred on later dates at higher elevations (Fig. 1). On average, the mean lapse rate for the onset of bud swelling was 3.1 ± 0.5 days/100 m in elevation gain; the difference in timing of bud swelling between the lowest and highest sites was 18 ± 3 days. Leaf unfolding began in late June and ended in mid-July; the mean lapse rate for this phenophase was 3.0 ± 0.6 days/100 m, leading to a difference in timing of 17 ± 3days between the lowest and highest sites. Although the lapse rate and duration of each phenophase did not differ among trees on the north- and southeast-facing sites, the mean dates of bud swelling and leaf unfolding were 2 ± 1 days later for trees growing on the north-facing sites than southeast-facing sites.

**Fig. 1.**
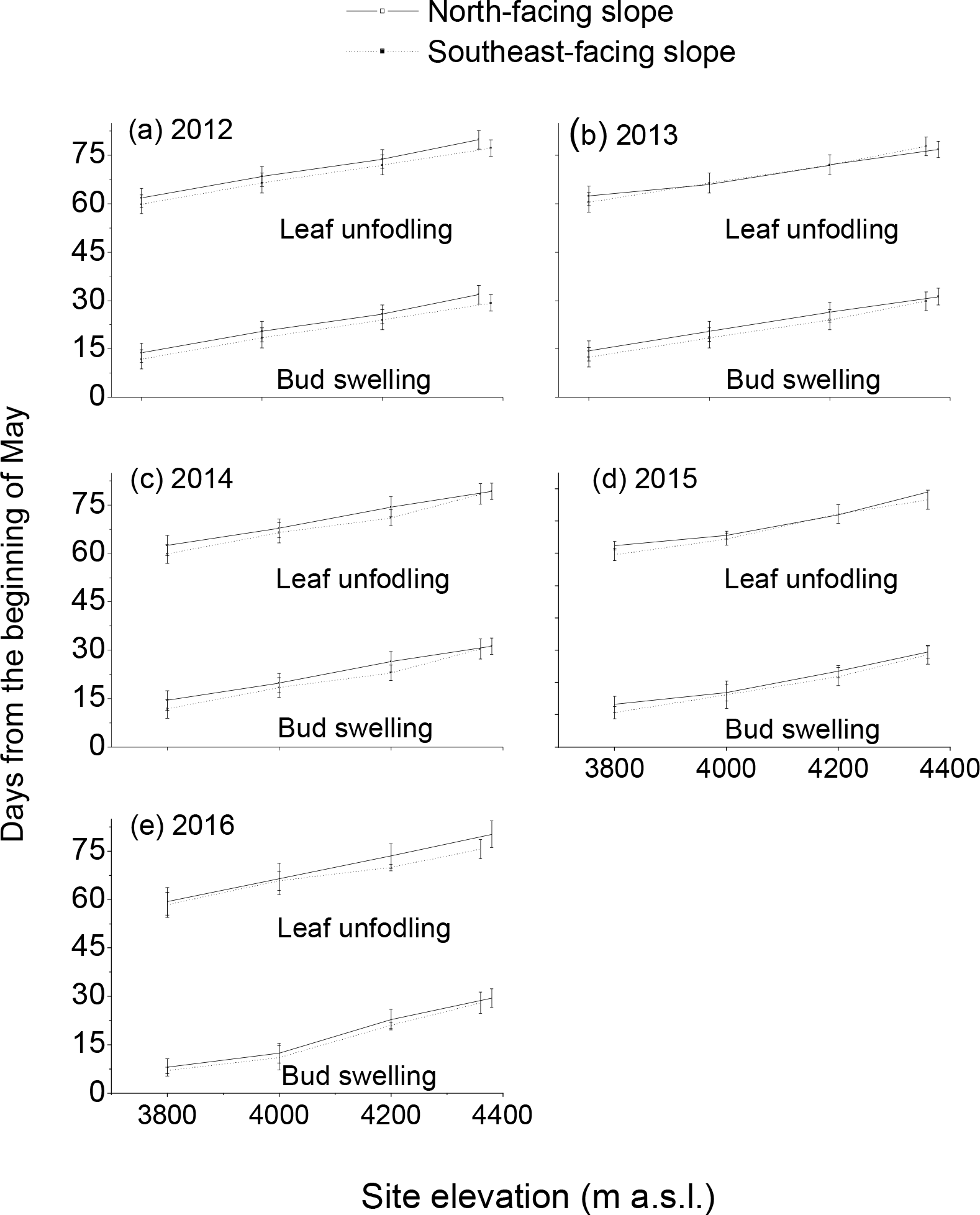
Variations (means ± SD) in onset of bud swelling and leaf unfolding with altitude and aspect during four study years.

### Explanatory modelling of phenophase timings

Our random forest models overall explained 84.8% of the variations in the onset of bud swelling and 93.1% of the variation in the onset of leaf unfolding. The frequency of spring freezing events and the last frost date were the most important predictors of bud swelling date (Fig. 2a). Of less importance were growing-degree days, maximum temperature, the minimum temperature, the measurement year, chilling days, and finally aspect (Fig. 2a). Mean safety margins were ≥ 7±5 days and increased significantly with increasing elevation (*r* = 0.24, *P* < 0.0001, *n* = 364) and among years (*r* = 0.27, *P* < 0.0001, *n* = 364), but not with aspect (*r* = 0.04, *P* = 0.36, *n* = 364). For leaf unfolding, onset of bud swelling was the most important predictor; the number of growing degree-days, minimum and maximum temperature ranked considerably lower, but were basically equivalent, in importance, followed by measurement year and aspect(Fig. 2b).

**Fig. 2.**
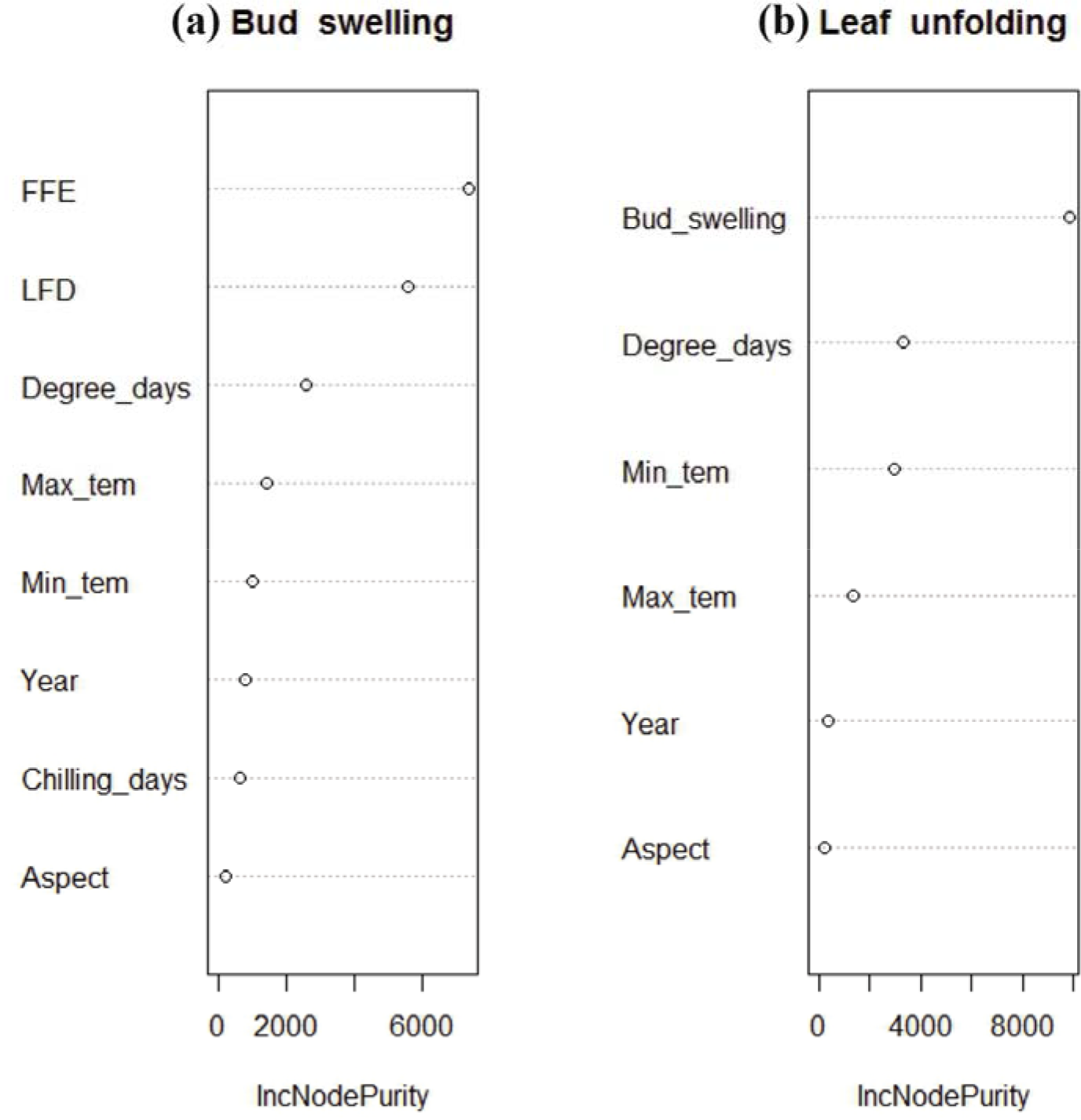
Ranked, relative importance of variables included in random forest models explaining variation in the onset dates of (a) bud swelling and (b) leaf unfolding.

Partial dependence plots indicated that the three response variables often were related non-linearly to the predictor variables (Fig. 3), which in turn interacted in complex ways (Fig. 4). Bud swelling was positively and essentially linearly related to both the frequency of spring freezing events and the last freezing date, negatively related to the sum of growing degree-days and minimum temperature (Fig. 3a), and non-linearly related to the other variables (Fig. 3a). Conditional inference tree modelling suggested that whether bud swelling occurred at later or earlier dates was largely controlled by the frequency of spring freezing events, with locations experiencing more than 62 freezing days in spring having the latest bud swelling dates; for earlier-onset locations (less than 62 freezing days), other factors such as last freezing date, the sum of growing degree daysinteracted to explain variability in earlier bud swelling timing (Fig. 4a).

**Fig. 3.**
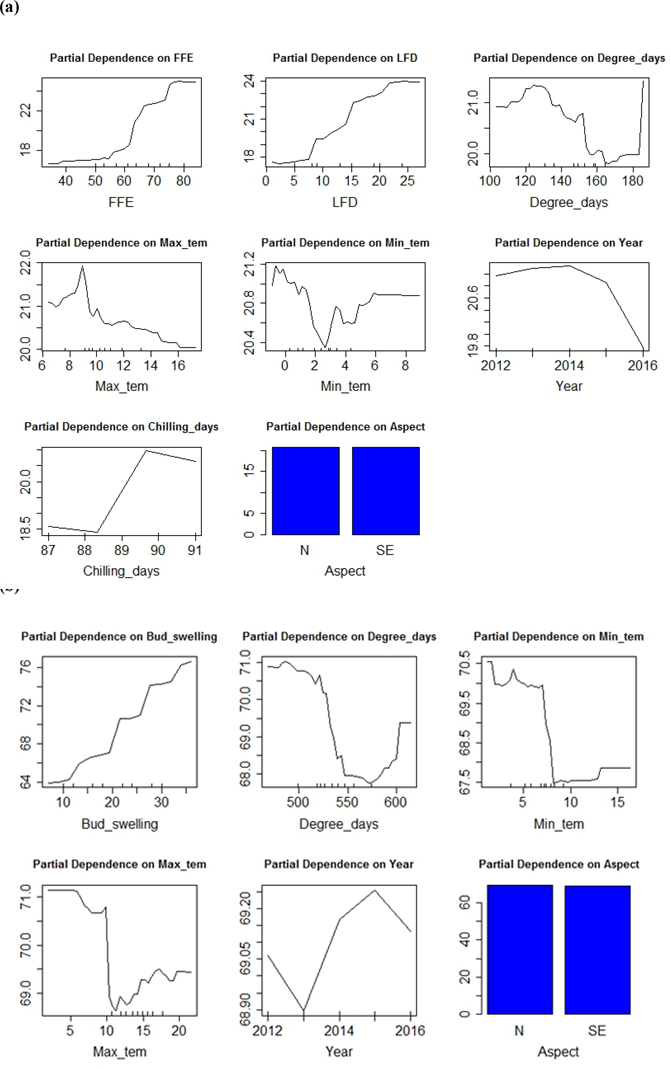
Partial dependence plots, based on results from the random forest analysis, showing the mean marginal influence of predictor variables on the onset date of (a) bud swelling and (b) leaf unfolding. Each plot represents the effect of one predictor variable on the response, while holding the other predictor variables constant at their mean values.

**Fig. 4.**
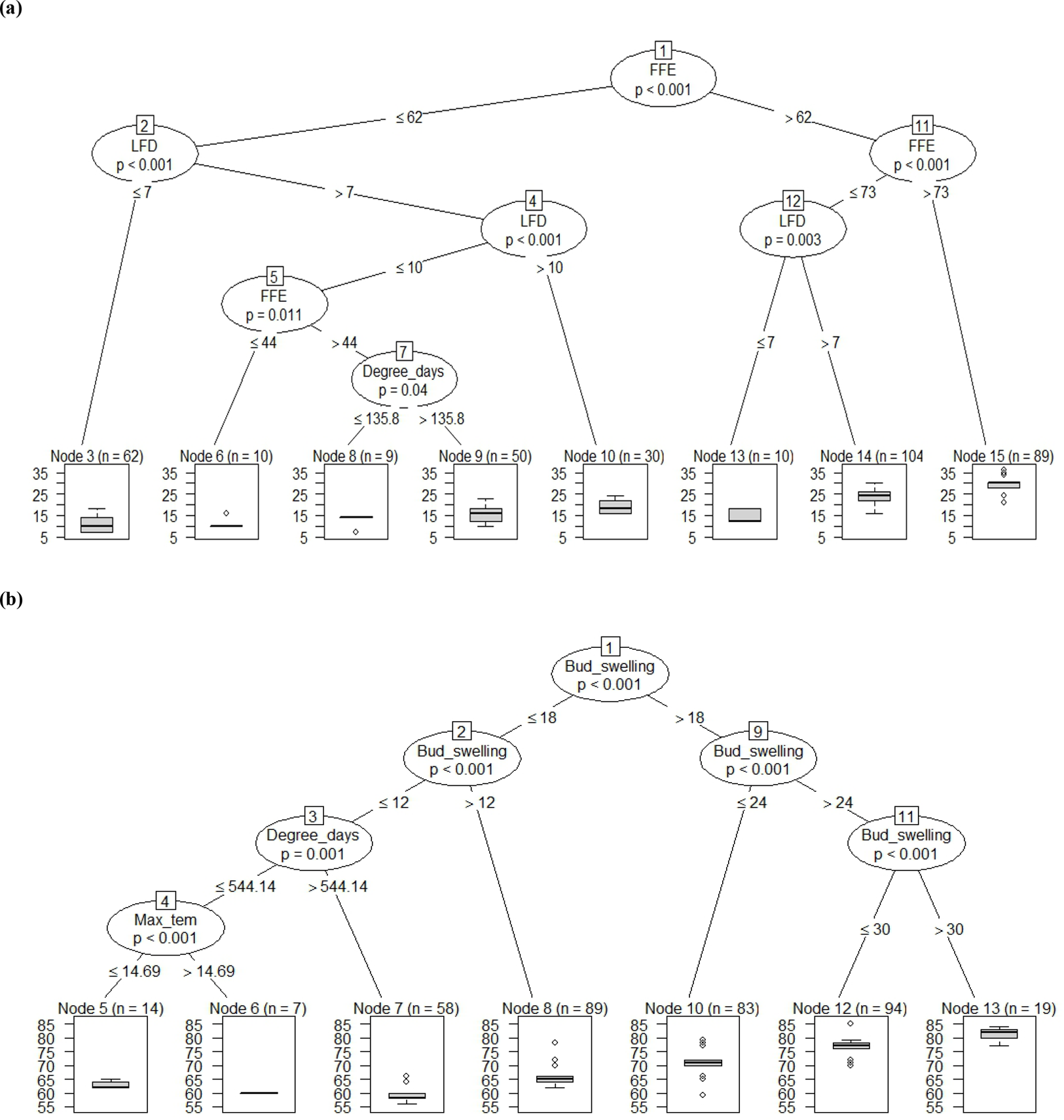
Conditional inference trees explaining variation in the onset date of: (a) bud swelling and (b) leaf unfolding based on sets of predictor variables (see Table 1). The trees show pathways of how the response data were recursively partitioned based on predictor variables. The observations associated with each terminal node are the result of these partitionings. P values at each node are from a Monte Carlo randomization test; in order for a split to occur P < 0.05.

The timing of leaf unfolding onset was positively and linearly related to onset of bud swelling, and had a negative, *s*-shaped relationship with growing degree days, with a switch from later to earlier timing of leaf unfolding past a critical value for the sum of growing degree-days that fell within the range of 530 – 540 accumulated °C (Fig 4b); a similar critical minimum temperatures of 7 – 7.5 °C was also evident (Fig. 3b). The relationship between leaf unfolding date and maximum temperature was more complex (Fig. 3b) likely due to interaction effects with the sum of growing degree days, maximum and minimum temperature in explaining finer variation in this phenophase (Fig. 4b).

## Discussion

A number of climatic variables can influence the onset and duration of plant phenophases, but research to date has tended to focus on temperature means rather than variance or extremes (Inouye, 2008; Wang et al., 2014), and on later phenophases such as flowering and leaf emergence. In contrast, the data we present here illustrates that timing of extreme events such as late spring frost and freezes, can strongly affect two phenophases, onset of bud swelling and leaf unfolding. The onset of bud swelling is of particular importance, because it is a prerequisite to the other important and well-studied phenophases. Furthermore, we took advantage of five years of data across a steep elevational gradient to provide additional information on climatic control of bud swelling and leaf unfolding.

The date of onset of bud swelling increased with elevation in all five years of this study. This result can be attributed directly to effects of temperature, as tree growth of Smith fir at high elevations is known to be limited by temperature (Liang *et al*., 2010; Wang *et al*., 2012). However, at high altitudes, trees are frequently exposed to large diurnal temperature fluctuations in spring (Ernakovich *et al*., 2014) and meristemetic tissues are especially vulnerable to damage from spring freezing (Gu *et al*., 2008). Once development starts in spring, freezing resistance is irreversibly lost and trees cannot re-acclimate to low temperatures (Lenz *et al*., 2013). In particular, the freezing resistance of trees decreases quickly as temperature increases during the de-hardening period (Lenz *et al*., 2013). In some cases, abnormally warm weather followed by sudden cold waves (particularly freezing events) in early to mid-spring can have disastrous impacts on plants effects (Gu *et al*., 2008). Thus, adaptations for avoidance of spring freezing damage are critical for the survival and subsequent development of many tree species in temperate and cold regions (Kollas *et al*., 2014). In our study, the frequency of freezing events and the date of the last hard freeze were the critical factors in regulating the timing of bud swelling. The normal date in late May of the last freeze that we observed matched that seen in the timing of snow melt at treeline in another study (Liu *et al*., 2011). Safety margin for spring frost increased significantly with elevation, suggesting the strong directional selection due to late spring frosts and freezes (Dantec *et al*., 2015). The negative effects of spring freezes on the survival of Smith fir seedlings (tree age ≤ 5yrs) had also been reported on the southeastern Tibetan Plateau (Shen *et al*., 2014). It is likely that Smith fir escapes from the spring freezing injury at the upper treelines by delaying bud swelling until very late spring (Wang *et al*., 2014).

Our results in combination with those from these other studies together support our hypothesis that frost-avoidance occurs in early phenophases of Smith fir, and our data provide the first empirical evidence that onset of spring phenology on the Tibetan Plateau is controlled primarily by spring frost, rather than the sum of growing degree-days.

Overall, these results are in line with phenological studies of deciduous broad-leaf trees in the eastern and western Alps (Lenz *et al*., 2013; Kollas *et al*., 2014) and Mount Fuji in Japan (Gansert, 2002), but different from what has been observed in warmer temperate forests where accumulation of growing degree-days is related more closely to phenological events (Peñuelas & Filella, 2001; Parmesan & Yohe, 2003; Wang *et al*., 2011; Dai *et al*., 2014). Our results partly explain the declining global warming effects on spring phenophases (Shen et al., 2013), because climatic extreme events (e.g. spring frosts) tend to increase with warming (Inouye, 2000; Augspurger, 2013; IPCC, 2013). Although the accumulation of chilling days has been shown to regulate the responses of spring phenology to climatic warming in Europe (Fu *et al*., 2015), the impact of chilling on the timing of bud swelling was negligible in our study. In addition, green-up date for grassland on the Tibetan Plateau is statistically related with mean minimum temperature during the preseason in both arid and wet regions (Shen *et al*., 2016). Comparisons between the findings of Shen et al. (2016) and our results suggest that spring phenology of different alpine plant function types might respond to different temperature variables.

The onset of leaf unfolding from the lowest to highest sites occurred in the warmest period of the growing season (from late June to mid-July) when freezing events were absent, and was driven more by the onset of bud swelling. This result suggested that earlier bud swelling translated into earlier leaf unfolding. Presumably, the annual growth cycle of trees forms an integrated system, where one phenophase can affect or regulate the subsequent phase (Fu *et al*., 2014a). Such carryover effects have also been reported in temperate and boreal forests (see review in Fu *et al*., 2014a). Unexpectedly, the sum of growing degree-days played a secondary role in controlling onset of leaf unfolding. These results are different from other studies conducted in temperate, boreal, and some alpine forests where accumulation of degree-days is the major determinant of the onset of leaf unfolding (Peñuelas & Filella, 2001; Parmesan & Yohe, 2003; Wang *et al*., 2011; Dai *et al*., 2014).

Last, our data provide evidence that warming may promote an advance in the timing of spring phenophase. The elevational transect that we studied represents a temperature gradient of ca. 3.8 °C, or a lapse rate of –0.66 °C/100 m (Liang *et al*., 2011). Therefore if the regional temperature warms by ≈ 1 °C, spring phenology could advance by 4.5 days, similar to the advancement of 4.6 days °C found in a meta-analysis of temperate plants around the world (Wolkovich *et al*., 2012). It appears that the temperature sensitivity of bud swelling and leaf unfolding phenology is somewhat greater than the sensitivity of flowering phenology which was found to be 3.3 days/ 1°C warming (Miller-Rushing & Primack, 2008). Earlier spring phenologies that accompany a warming climate also may help to explain some observed upward shifts in alpine treelines, The 1.2 – 1.5 °C warming observed on the Tibetan Plateau in the last 100 years has been paralleled by an up to 80-m upslope shift in treeline if species interactions do not constrain these shifts (Liang *et al*., 2016).

## Acknowledgements

This work was supported by the National Natural Science Foundation of China (41525001, 41661144040) and the International Partnership Program of Chinese Academy of Sciences (131C11KYSB20160061). AME’s participation in this project was supported by the Chinese Academy of Sciences (CAS) President International Fellowship Initiative for Visiting Scientists, Grant no. 2016VBA074. We appreciate the great support from the Southeast Tibet Station for Alpine Environment, Observation and Research, CAS.

## Supporting information captions

**Table S1.**
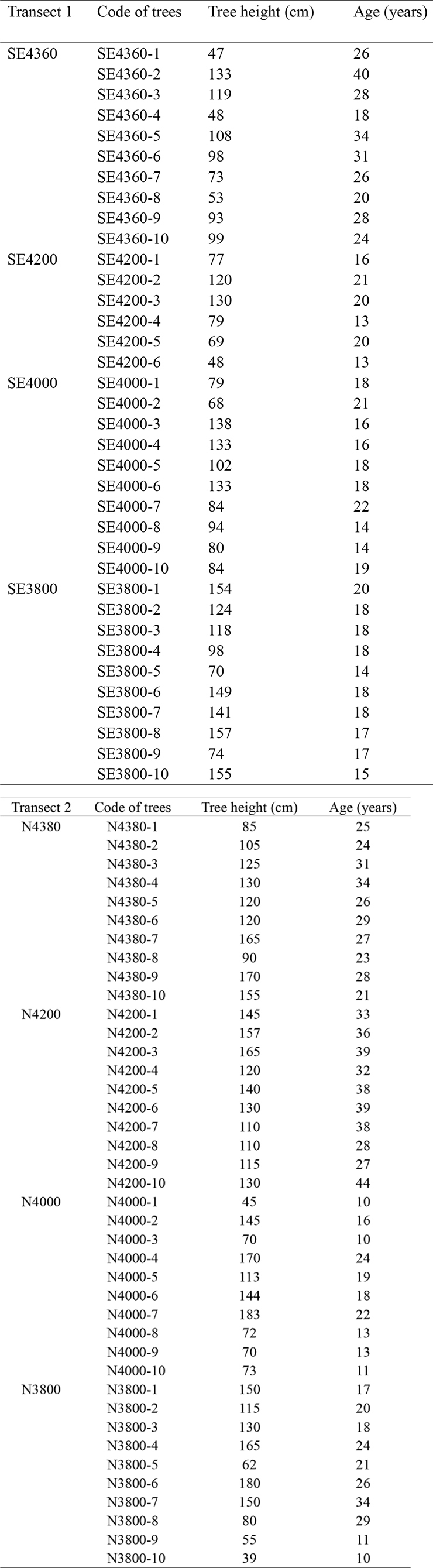
*Tree height and age of Smith fir individuals at four elevations along two altitudinal transects. Note that the height and age of each sampled tree was determined in April 2012, when buds were dormant.*

**Fig. S1.**
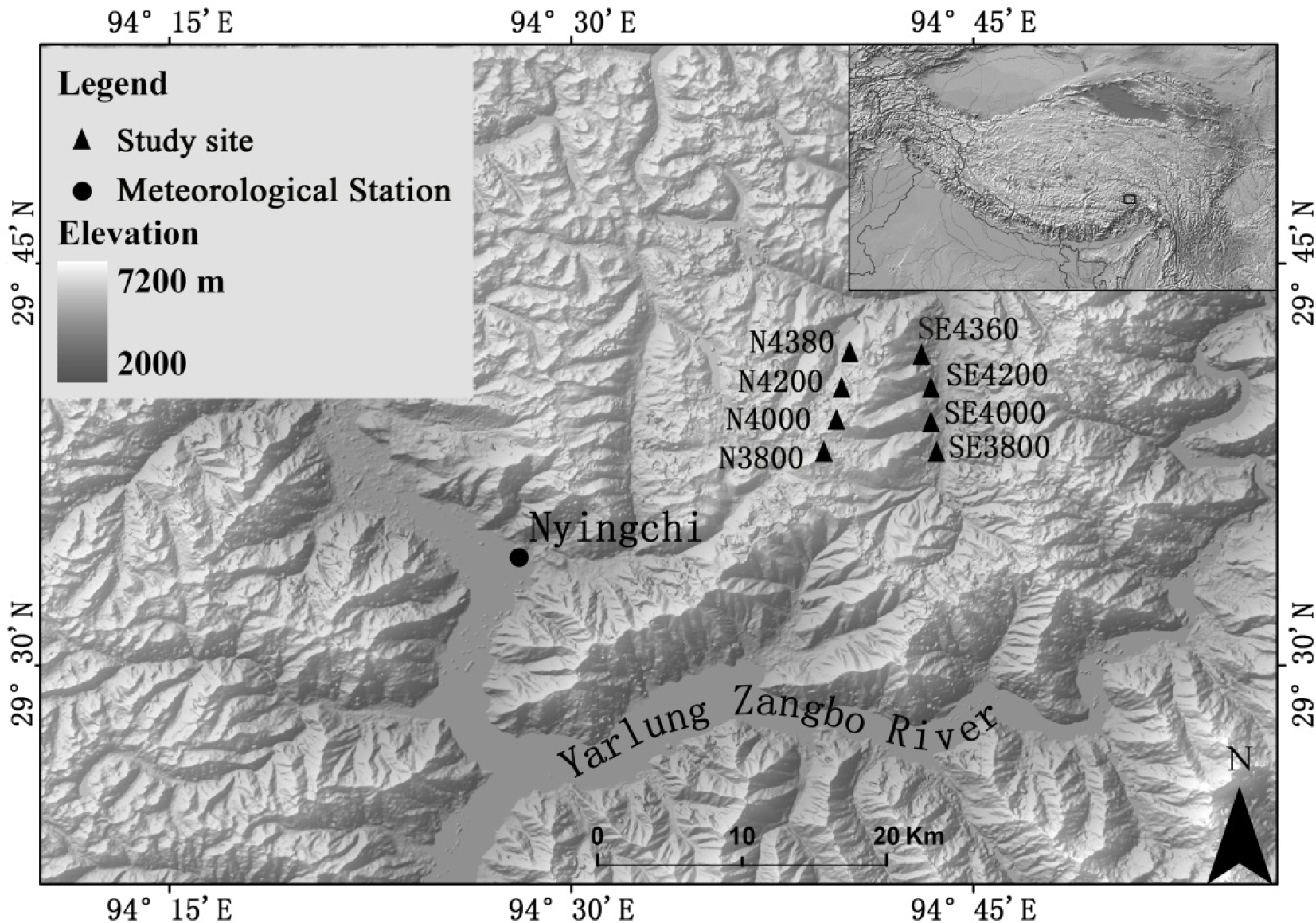
Location of the study sites, the meteorological station at Nyingchi (3,000 m a.s.l.) in the southeastern Tibet. SE andN and the number represent the slopes and the site elevation (m a.s.l.), respectively. Inset (upper right corner) indicates the position of the study region on the Tibetan Plateau.

